# Quantum Barcoding Bridges High-Dimensional Immune Phenotyping with Signaling Responses across Species

**DOI:** 10.64898/2026.10.31.763237

**Authors:** Tze-Kai Tan, Kevin Taylor, Fan Zhang, Yunhao Bai, Bokai Zhu, Marc A. Baertsch, Sasha Anronikov, Zachary Bjornson-Hooper, Debbie L. L. Ho, Angelica Trejo, Hiromitsu Nakauchi, Agnieszka Czechowicz, Frank Steemers, Garry P. Nolan, David R. McIlwain

**Author notes:** Senior authors (Hiromitsu Nakauchi, Agnieszka Czechowicz, Frank Steemers, Garry P. Nolan, David R. McIlwain). Lead contact: David R. McIlwain.

## Abstract

Cross-species immune profiling is limited by marker-panel size and antibody cross-reactivity. We benchmark Quantum Barcoding (QBC), a split-pool oligo-antibody assay, against mass cytometry in human and rhesus macaque blood and integrate separately measured cells using MARIO. The resulting computationally matched profiles carry 174 protein features. Population-frequency concordance was higher in humans (Pearson r = 0.92) than rhesus macaques (r = 0.55), and fusion resolved complementary lineage subsets. Unstimulated signaling conservation varied by lineage, with B-cell profiles correlated across species (Spearman ρ = 0.70). In the primary PBMC-reference analysis, GM-CSF increased pSTAT5 in human monocytes and dendritic populations and rhesus dendritic cells. Network inference identified convergent pSTAT5-centered connectivity changes in human monocytes and rhesus dendritic cells, including pSTAT1-edge reversal and gained pSTAT3/pERK coupling. These results connect broad surface phenotypes to inferred signaling organization while retaining species- and population-specific differences.

**In Brief (eTOC blurb):** Tan and colleagues computationally match Quantum Barcoding and mass-cytometry profiles from human and rhesus macaque immune cells. The 174-feature resource resolves complementary cell populations and links GM-CSF-induced pSTAT5 responses to convergent inferred network changes in human monocytes and rhesus dendritic cells.

**Highlights:**

- Computational QBC–CyTOF matching links 174 features in paired cell profiles
- Population-frequency concordance is higher in human than rhesus PBMCs
- GM-CSF increases pSTAT5 in human monocytes and rhesus dendritic cells
- Inferred pSTAT5 neighborhoods show convergent rewiring across species

## Introduction

Single-cell technologies have transformed the analysis of immune-cell heterogeneity, but protein-level profiling remains constrained by marker-panel size. Mass cytometry (CyTOF) measures surface and intracellular proteins using metal-isotope-tagged antibodies^1,2^. Advances in metal conjugation and barcoding have expanded its measurement and multiplexing capabilities^3,4,5,6,7,8^. Sequencing-based antibody tags provide an alternative readout, exemplified by CITE-seq^9^. NEAT-seq, QuRIE-seq and Phospho-seq extend sequencing-based assays to nuclear, intracellular or phosphorylated proteins^10,11,12^. Combining broad surface phenotyping with established intracellular signaling panels remains an important objective for comparative immunology.

Split-pool combinatorial synthesis generates cell barcodes through iterative oligonucleotide indexing rather than isolation into individual wells or droplets^13,14,15^. Quantum Barcoding (QBC) applies this principle to antibody–oligonucleotide conjugates. Its original demonstration measured up to 50 targeted proteins and, separately, more than 70 RNA regions, and illustrated joint protein/RNA profiling ^14^. Here we use a 134-marker QBC surface-antibody panel and computationally connect it to CyTOF intracellular signaling measurements.

Cross-species translational immunology provides a setting in which these complementary measurements are useful. Rhesus macaques are important non-human primate models for studying human immune biology^16,17^. Comparative profiling has documented species-specific differences in immune-cell composition, unstimulated signaling and stimulus responses^18,19^. Limited cross-reactivity of human-optimized antibody reagents further complicates direct comparison. Methods that connect surface identity with intracellular signaling can help identify both shared responses and population-specific differences without assuming cross-species equivalence.

Here we benchmark QBC against CyTOF in three human and three rhesus macaque donors, then integrate the modalities using MARIO, a partial-overlap cell-matching algorithm^20^. Each resulting profile links separately measured QBC and CyTOF cells through shared surface markers and carries 174 features; it is not a same-physical-cell measurement. We also match the unstimulated QBC reference to a published CyTOF cross-species stimulation dataset^18^ to ask which signaling responses are shared, which vary by population or species, and how inferred signaling connectivity changes with stimulation. The primary PBMC-reference analysis identifies GM-CSF-induced pSTAT5 responses in human monocytes and dendritic populations and rhesus dendritic cells. Network analysis reveals convergent pSTAT5-centered neighborhood changes in human monocytes and rhesus dendritic cells, alongside heterogeneous unstimulated and stimulus-response profiles.

## Results

### QBC and CyTOF provide complementary immune lineage detection resolution

Quantum Barcoding (QBC) adds combinatorial DNA indices to antibody–oligonucleotide conjugates through split-pool synthesis, with molecular counts obtained from unique molecular identifiers (UMIs) (Figure 1A, 1B; STAR Methods). Four 60-way split-pool barcode rounds were used, corresponding to 60^4^ = 12,960,000 nominal combinations. These are combinatorial capacities, not measured singlet yields or collision rates. Library amplification used 50,000 cells per PCR (STAR Methods; Figure S1).

**Figure 1.**
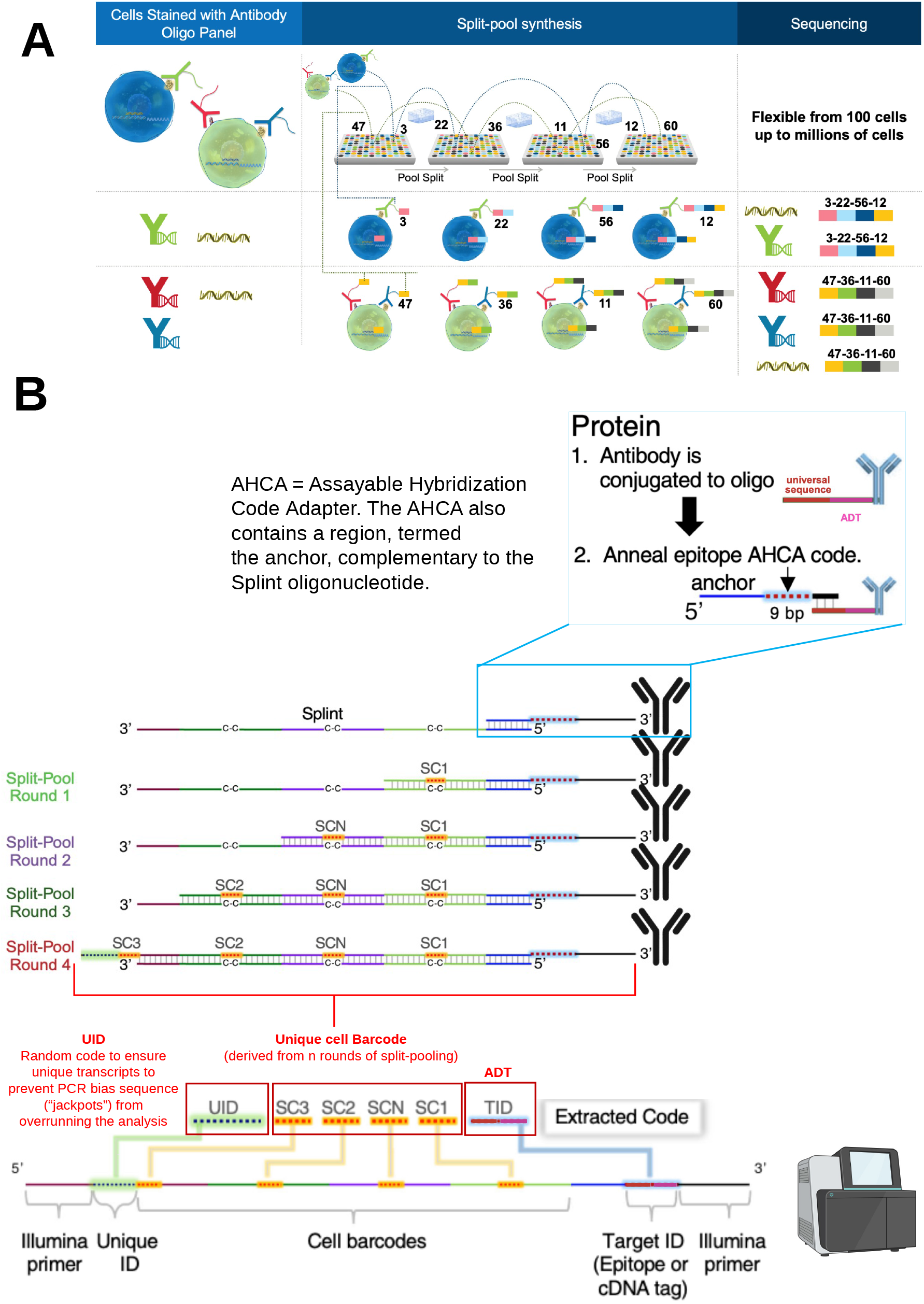
Split-pool Single-Cell Quantum Barcoding (QBC) Methodology and Schematics to enable High-Throughput Single-Cell Proteomic Profiling. (A) Overall scheme of split-pool single-cell quantum barcoding. Immunostained cells carrying antibody– oligonucleotide conjugates are barcoded through iterative split-pool synthesis. The implemented workflow used four 60-way split-pool rounds (SC-A through SC-D), giving a nominal capacity of 60^4^ = 12,960,000 cell-barcode combinations (STAR Methods). (B) AHCA (Assayable Hybridization Code Adapter) schematic. The AHCA contains an anchor region complementary to the Splint oligonucleotide, a unique cell-barcode region derived from n rounds of split-pooling, a UID (random code to ensure unique transcripts and prevent PCR jackpot bias), and an antibody-derived tag (ADT) region. Schematic adapted from O’Huallachain et al., 2020^14^.

To validate sample barcoding, we pooled two human PBMC samples each pre-stained with a distinct hashtag and ran the TBNK panel through the QBC workflow (Figure 2A). A UMI–rank knee plot established a 1,000-UMI cutoff and yielded 1,327 high-quality cells, cleanly separating CD3⁺ T cells, CD3⁺CD8⁺ T cells, and CD3⁻ non-T cells across the two hashtag-resolved samples (Figure 2B).

**Figure 2.**
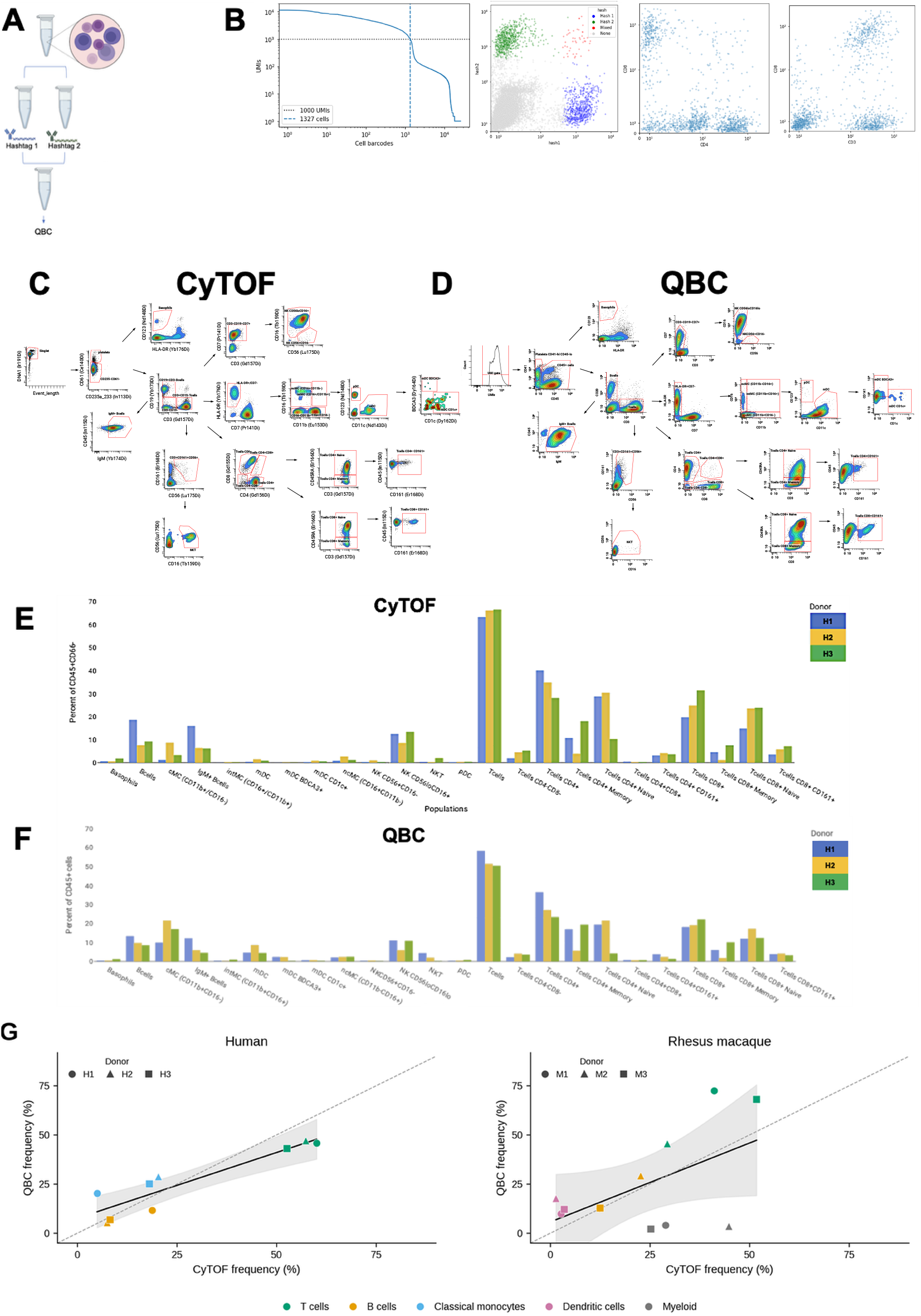
QBC–CyTOF population-frequency benchmarking. (A) Validation of QBC using hashtag antibodies. Barnyard experiment using human cells with the BioLegend TBNK panel and two-level split-pool barcoding. (B) Plot displays the total number of Unique Molecular Identifiers (UMIs) for each barcode, ranked in descending order. This “knee plot” is used to distinguish barcodes associated with viable cells (plateau region) from those representing empty events (steep drop-off). A UMI count threshold of 1,000 UMIs (horizontal dotted line) was applied to filter the data, resulting in the selection of 1,327 high-quality cells (indicated by the vertical dashed line) for downstream analysis. Right panels: hashtag-resolved scatter plots (Hash1 vs Hash2; CD4 vs CD3; CD8 vs CD3) confirm sample-of-origin assignment and T-cell subset detection. (C, D) Manual gating of CyTOF (C) and QBC (D) datasets identifies major immune cell populations using CyTOF and QBC panels with matched major immune cell markers (representative sample, donor H1). (E, F) Population frequencies for each gated population across the three human donors (H1–H3) measured by CyTOF (E; percent of CD45⁺CD66⁻ cells) and QBC (F; percent of CD45⁺ cells), showing concordant major-lineage composition and donor-specific signatures (elevated basophils in H1, B cells in H2, naïve CD8⁺ T cells in H3). (G) Population frequencies measured independently by QBC (y-axis) and CyTOF (x-axis). Each point is one canonical population in one donor: human B cells, T cells and classical monocytes (left; n = 9 population–donor pairs), or rhesus B cells, dendritic cells, myeloid cells and T cells (right; n = 12 pairs). Color denotes population and shape denotes donor (H1–H3 or M1–M3). The original saved frequency values are unchanged. The solid line is an ordinary least-squares fit, the gray band its nominal point-level 95% confidence interval, and the dashed line equality. Human Pearson r = 0.917, RZ = 0.841, P = 5.0 × 10^−4^; rhesus r = 0.553, RZ = 0.306, P = 0.062. There are three donors per species; repeated populations within donors are not independent biological replicates.

We benchmarked QBC against CyTOF on cryopreserved PBMCs from three human donors (H1–H3) and three rhesus macaque donors (M1–M3). The study-specific CyTOF panels contained 42 human and 43 rhesus surface and intracellular markers (Table S1), using a workflow adapted from published cross-species profiling^18,19^. QBC used the 134-marker BioLegend TotalSeq-B Universal v1.0 antibody cocktail (BioLegend; Table S2); 18 markers were shared with CyTOF (Table S3). Manual gating identified major immune lineages in both modalities, including platelet, basophil, T-cell, B-cell, NK/NKT, dendritic and monocyte compartments (Figure 2C, 2D). Donor-specific composition differences included the elevated basophil fraction in H1, B-cell fraction in H2 and naïve CD8 T-cell fraction in H3 (Figure 2E, 2F).

Relative frequencies of harmonized canonical populations were correlated across platforms in human PBMCs (Pearson r = 0.92, R^2^ = 0.84, P = 4.96 × 10^-4^; n = 9 population–donor pairs from three donors; Figure 2G, left). This is a descriptive population-frequency benchmark, not a direct assessment of per-cell matching accuracy. The nominal point-level test does not account for within-donor dependence or compositional constraints.

In rhesus macaques, cross-platform concordance was moderate and did not reach significance (Pearson r = 0.55, R^2^ = 0.306, P = 0.062; n = 12 population–donor pairs from three animals; Figure 2G, right). This is consistent with limitations of a human-optimized QBC antibody cocktail in rhesus cells, although population definitions and gating also affect frequency agreement. We therefore used a curated 59-marker QBC subset as a common cross-species feature space for fused clustering (Table S4).

### MARIO fusion of QBC and CyTOF dataset overcomes modality-specific blind spots in human and rhesus PBMCs

QBC provides broad surface phenotyping, whereas CyTOF includes an established intracellular signaling panel. We computationally linked separately measured cells from the paired unstimulated datasets, generated from the same three donors per species, using MARIO^20^ and 18 shared lineage markers (Table S3). The matched profiles carry 174 measurements—134 QBC and 40 CyTOF features—combining complementary information without asserting physical same-cell co-measurement.

We used the curated 59-marker QBC subset for cross-species clustering of the fused resource (Table S4). The standalone unstimulated datasets were analyzed separately using their saved modality-specific feature selections and cluster annotations (Figure 3A–D; STAR Methods). Human CyTOF-only clustering resolved ten annotated populations, including conventional and plasmacytoid dendritic cells and MAIT cells, but did not separately resolve NK cells, platelets or non-classical monocytes under these settings (Figure 3A). Human QBC-only clustering resolved seven populations, including NK cells, platelets and γδ T cells, but not dendritic or non-classical monocyte populations (Figure 3C). Rhesus QBC-only clustering resolved six populations and left 64.7% of cells unassigned; it did not separately resolve a CD4 T-cell population (Figure 3D). Rhesus CyTOF-only clustering resolved nine populations, including NK cells and dendritic and granulocyte-containing populations, but no platelet population (Figure 3B). These differences describe the displayed analyses, not absolute detection limits of either platform.

**Figure 3.**
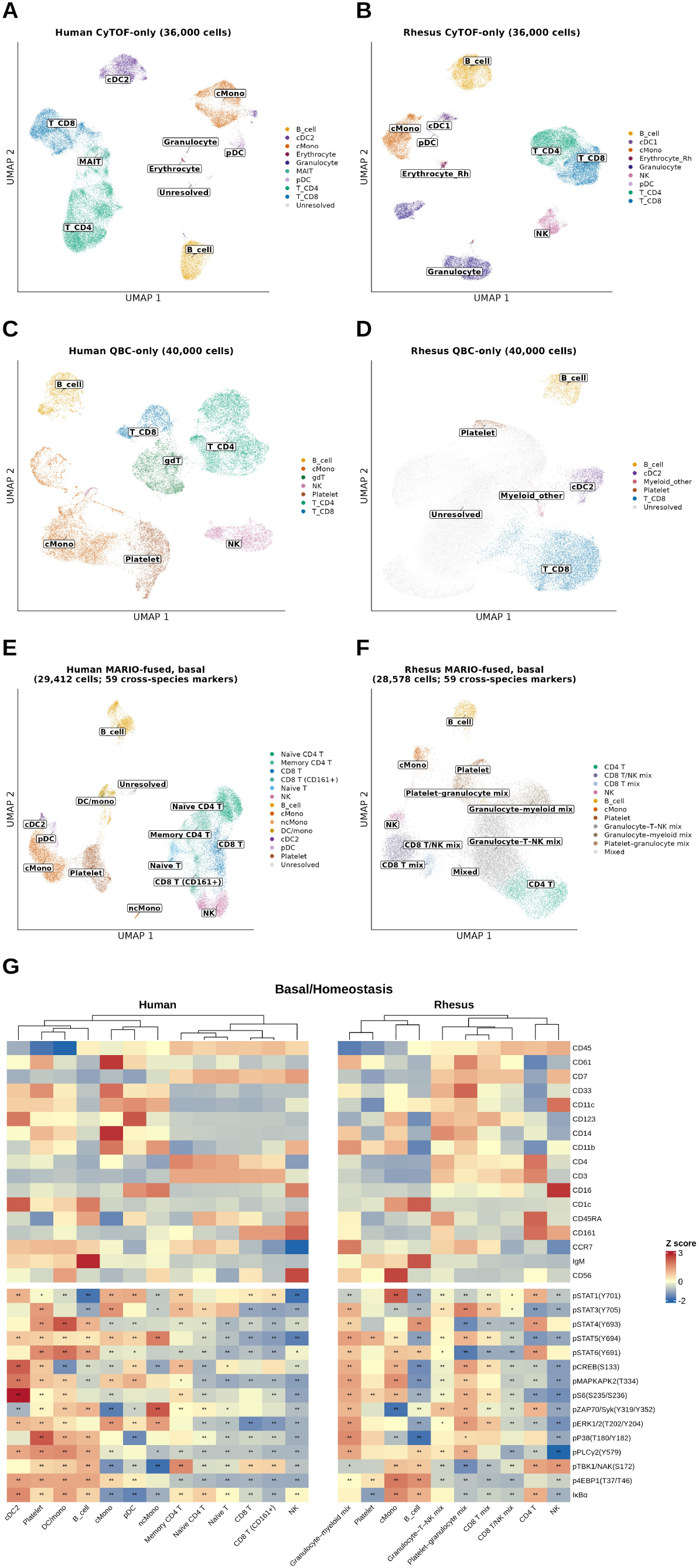
Complementary population resolution in unstimulated QBC– CyTOF fusion. (A–D) UMAP views of standalone unstimulated datasets, colored by saved population annotations. PCA feature sets and selected resolution-0.3 assignments are described in STAR Methods. (A) Human CyTOF-only: 36,000 cells and ten annotated populations. (B) Rhesus CyTOF-only: 36,000 cells and nine populations. (C) Human QBC-only: 40,000 cells and seven populations. (D) Rhesus QBC-only: 40,000 cells and six populations, with 25,889/40,000 cells unassigned (64.7%). CyTOF includes 12,000 cells per donor. QBC includes 10,000 Stanford cells per donor plus 10,000 additional Davis cells from H1 or M3 in the respective species, not an additional donor. (E, F) Unstimulated MARIO-matched profiles clustered in the 59-marker QBC space (Table S4) and colored by the final saved annotation map. Human profiles comprise 29,412 pairs with fourteen annotated populations; rhesus profiles comprise 28,578 pairs with eleven annotated populations. These labels consolidate 18 human and 13 rhesus graph clusters and are distinct from stimulation clustering in Figure 4. QBC-side donor contributions are shown in Figure S2. Mixed rhesus labels denote unresolved source-annotation mixtures, not uniquely identified granulocyte QBC counterparts. (G) Unstimulated signaling states of annotated populations in the MARIO-fused unstimulated datasets. Heatmaps show row-wise z-scored population means for 17 surface context markers and 15 intracellular signaling nodes in human (left) and rhesus (right); columns are hierarchically clustered within species (Pearson distance, average linkage). Asterisks mark signaling nodes where a population deviates significantly from all other cells (donor-adjusted linear models, BH-FDR: * padj < 0.05, ** padj < 0.01). Cross-species conservation per shared lineage was assessed by Spearman correlation of 15-node profiles with permutation tests (10,000 shuffles, BH-FDR); node-level species differences used donor-aware mixed-effects models. Signaling values derive from the fused dataset and are compressed relative to raw arcsinh-transformed CyTOF values; relative patterns across populations are interpretable.

Clustering the unstimulated fused datasets in the 59-marker space resolved complementary populations in both species (Figure 3E, 3F). The final annotation map consolidated the human graph clusters into fourteen populations, including NK cells, platelets, non-classical monocytes, plasmacytoid and conventional DC2 populations, naïve/memory CD4 T cells and multiple CD8 T-cell populations. The rhesus map yielded eleven annotated compartments, recovering B cells, CD4 T cells, NK cells, classical monocytes and platelets from the QBC-unresolved fraction. Some rhesus macaque clusters retained mixed labels because the 59-marker panel did not distinguish granulocyte-containing source annotations from other lineages; dedicated granulocyte markers would be needed to resolve these mixtures. Donor-colored embeddings show contributions from multiple donors but do not establish absence of donor or site effects (Figure S2).

We examined unstimulated intracellular signaling across the fused annotations (Figure 3G). Myeloid, dendritic and platelet compartments showed elevated readouts across mTORC1, MAPK and pCREB nodes relative to quieter T-cell populations in the saved donor-adjusted analysis. Cross-species 15-node profiles showed modest global conservation (Spearman ρ = 0.37; permutation-adjusted P = 0.0098). B cells were the most conserved lineage (ρ = 0.70; adjusted P = 0.0098); classical monocytes showed an intermediate correlation (ρ = 0.36), while CD4 T, CD8 T, NK and platelet profiles were not significantly conserved. Rhesus cells had lower unstimulated total IκBα in B-cell, CD4 T-cell, NK and platelet compartments, together with elevated pSTAT5 (Y694) and pSTAT1 (Y701) in lymphoid lineages; human cells showed higher pTBK1/NAK (S172) and pCREB (S133). The IκBα measurement reports total protein abundance, not IκBα phosphorylation, and its direction alone does not establish NF-κB activation.

### Cross-modal stimulation fusion identifies a species-conserved GM-CSF– pSTAT5 axis

To examine stimulation-associated signaling, we matched the unstimulated QBC reference to unstimulated, CD40L and GM-CSF conditions from a published whole-blood CyTOF dataset comprising 46 human and 15 rhesus donors^18^. For the primary analysis, reference cells annotated as granulocyte/myeloid, platelet, erythrocyte or unresolved were excluded before matching because these compartments were not represented comparably in the PBMC QBC reference. Matching against the full whole-blood reference was retained as sensitivity analysis. The primary fused objects contained 20,862 human pairs (7,740 unstimulated, 6,164 CD40L, 6,958 GM-CSF) and 14,414 rhesus pairs (4,807, 4,618, 4,989; Figure 4A–4C). Matching assumes that shared surface phenotypes remain informative across the reference conditions; the QBC cells themselves were not stimulated. Conditions mixed extensively in the embeddings (median condition cLISI 2.5 and 2.7, maximum 3; Figure 4F, 4G). Each pair retained its CyTOF partner’s source population annotation; broad source labels define the eight human and six rhesus populations displayed in Figure 4D and 4E. Cluster-plurality annotations are reported separately as an audit (Table S5), not used to relabel these results.

**Figure 4.**
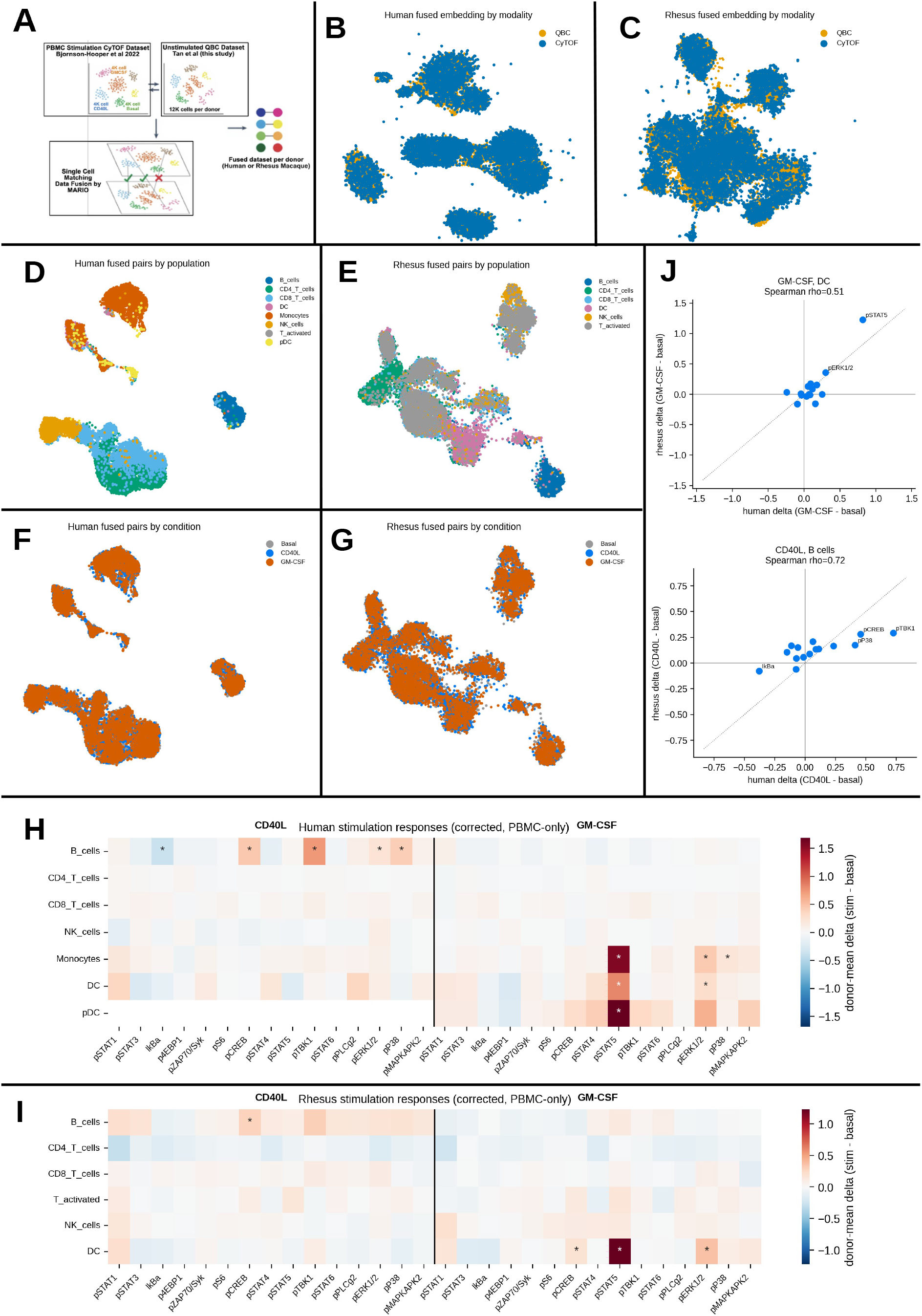
Cross-modal stimulation response in the MARIO fused QBC-CyTOF reference. (A) Computational workflow: unstimulated QBC surface profiles are linked through 18 shared markers to a PBMC-restricted CyTOF reference containing unstimulated, CD40L and GM-CSF conditions^18,20^. Each 174-feature profile represents a computational pair of separately measured cells. (B, C) Joint UMAP embeddings of matched QBC and CyTOF cells for human (B; 20,862 matched pairs) and rhesus macaque (C; 14,414 matched pairs), colored by modality (QBC, orange; CyTOF, blue). (D, E) Primary PBMC-reference fused-pair embeddings for human (D) and rhesus macaque (E), colored by broad population labels from the CyTOF source-cell partner. Graph clustering used 59 QBC features at resolution 0.5 (16 human and 11 rhesus clusters), but the displayed analysis labels were not generated by cluster-plurality assignment. Table S5 separately reports cluster composition, plurality-audit labels, purity and within-pair anchor correlation. (F, G) The same fused-pair embeddings colored by stimulation condition: unstimulated, CD40L, or GM-CSF. Median condition local inverse Simpson index (cLISI), calculated on the joint MARIO embedding, was 2.5 in human and 2.7 in rhesus macaque (maximum possible value, 3). (H, I) Stimulation-response heatmaps for human (H) and rhesus macaque (I), showing donor-mean changes relative to unstimulated (Δ = stimulated − unstimulated) across 15 intracellular signaling nodes and the indicated populations. CD40L and GM-CSF responses are displayed in separate blocks. Significance was assessed using paired, two-sided donor-level t-tests with Benjamini–Hochberg false-discovery-rate adjustment (adjusted P < 0.05, asterisk). The source dataset comprised 46 human and 15 rhesus macaque donors; individual tests required at least five paired donors, with at least three cells per donor in each compared condition. Color scales are species-specific: human, ±1.7; rhesus macaque, ±1.3. (J) Primary PBMC-reference cross-species response comparisons for GM-CSF in dendritic cells (top; Spearman ρ = 0.511; permutation P = 0.0553) and CD40L in B cells (bottom; ρ = 0.725; P = 0.00330). Each point is one of 15 intracellular signaling measurements, plotted by its human and rhesus donor-mean stimulated-minus-unstimulated response. These are not the whole-blood granulocyte sensitivity results. Dotted lines indicate equal responses; selected nodes are labeled. Permutation tests used 10,000 permutations.

We assessed matching using three comparisons (STAR Methods). Within-pair correlation across 18 shared anchors exceeded shuffled-partner correlations (human median r = 0.79 versus 0.09, with 88% of pairs above r = 0.5; rhesus median r = 0.43 versus 0.14). Harmony integration^21^ followed by nearest-neighbor label transfer agreed with MARIO source labels on 90.1% of human cells (adjusted Rand index, ARI = 0.83). CellFuse (preprint)^22^, a supervised classifier trained on the annotated CyTOF reference, agreed on 91.0% (ARI = 0.81). Rhesus agreement was lower at fine T-subset resolution and improved after T-subset collapse. These anchor- and reference-dependent comparisons assess consistency; they do not independently validate all imputed feature combinations.

In the primary PBMC-reference analysis, human GM-CSF responses were concentrated in myeloid populations: pSTAT5 (Y694) increased in monocytes (donor-mean Δ = +1.56; adjusted P = 3.0 × 10⁻Z⁶), plasmacytoid dendritic cells (+1.69; adjusted P = 4.6 × 10⁻3) and dendritic cells (+0.82; adjusted P = 1.7 × 10⁻3; Figure 4H). Monocyte responses also included pERK1/2 (+0.44; adjusted P = 2.8 × 10⁻⁷) and pP38 (+0.21; adjusted P = 1.3 × 10⁻3). No lymphoid response reached significance under the same testing criteria. CD40L elicited a B-cell program including pTBK1/NAK (+0.72; adjusted P = 3.0 × 10⁻¹¹), pCREB (+0.46; adjusted P = 2.8 × 10⁻⁷), pP38 (+0.41; adjusted P = 6.4 × 10⁻⁷), reduced total IκBα (−0.38; adjusted P = 3.4 × 10⁻⁵) and pERK1/2 (+0.23; adjusted P = 0.043).

In the primary rhesus analysis, GM-CSF increased dendritic-cell pSTAT5 (+1.23; adjusted P = 2.7 × 10⁻⁵), pERK1/2 (+0.36; adjusted P = 0.010) and pCREB (+0.17; adjusted P = 0.019; Figure 4I). CD40L increased B-cell pCREB (+0.28; adjusted P = 7.0 × 10⁻3). The separate whole-blood sensitivity analysis identified granulocyte/myeloid pSTAT5 increases in both species (human +1.50, adjusted P = 8.1 × 10⁻Z⁶; rhesus +1.29, adjusted P = 5.5 × 10⁻⁷). Because granulocytes lack a comparable QBC PBMC counterpart, those sensitivity results do not establish a matched granulocyte surface/signaling profile.

Cross-species concordance of the full 15-node response vector depended on stimulus and lineage (Figure 4J). In the primary analysis, the GM-CSF dendritic-cell correlation was positive but did not meet P < 0.05 (Spearman ρ = 0.511; permutation P = 0.0553). CD40L profiles were concordant in B cells (ρ = 0.725; P = 0.00330) and dendritic cells (ρ = 0.657; P = 0.00840). Across all shared lineages, concordance was significant for CD40L (ρ = 0.350; P = 0.000100) but not GM-CSF (ρ = 0.189; P = 0.264). The whole-blood sensitivity analysis gave a stronger GM-CSF granulocyte/myeloid correlation (ρ = 0.854; P = 0.000300) and CD40L B-cell correlation of 0.761 (P = 0.000800). Thus pSTAT5 induction occurs in myeloid populations of both species, but this does not imply significant conservation of the entire primary GM-CSF response vector.

### Score-rooted Pseudotime analysis resolves T-cell-linked surface-markers upon stimulation

We fitted Monocle3^31^ trajectories separately for each species and condition, generating four graphs for unstimulated and GM-CSF reference groups and excluding CD40L (Figure 5A–5D). Each graph used all 174 fused measurements. The QBC surface-phenotype reference comprised three unstimulated donors per species; condition labels came from the CyTOF partners. Graphs were rooted using the top 5% of rows by the nine-marker activation/proliferation score (five CyTOF and four QBC measurements; STAR Methods), whose values are displayed in Figure 5E and 5F. Because the score sets the roots, its association with pseudotime is not independent validation of progression. Given that pSTAT5 has the most significant change upon GM-CSF stimulation in human Classical Monocytes and *rhesus macaque* Plasmacytoid dendritic Cells and Dendritic Cells populations, population changes were compared as paired reference-donor GM-CSF-minus-unstimulated score differences, not as distances between independent pseudotime axes.

**Figure 5.**
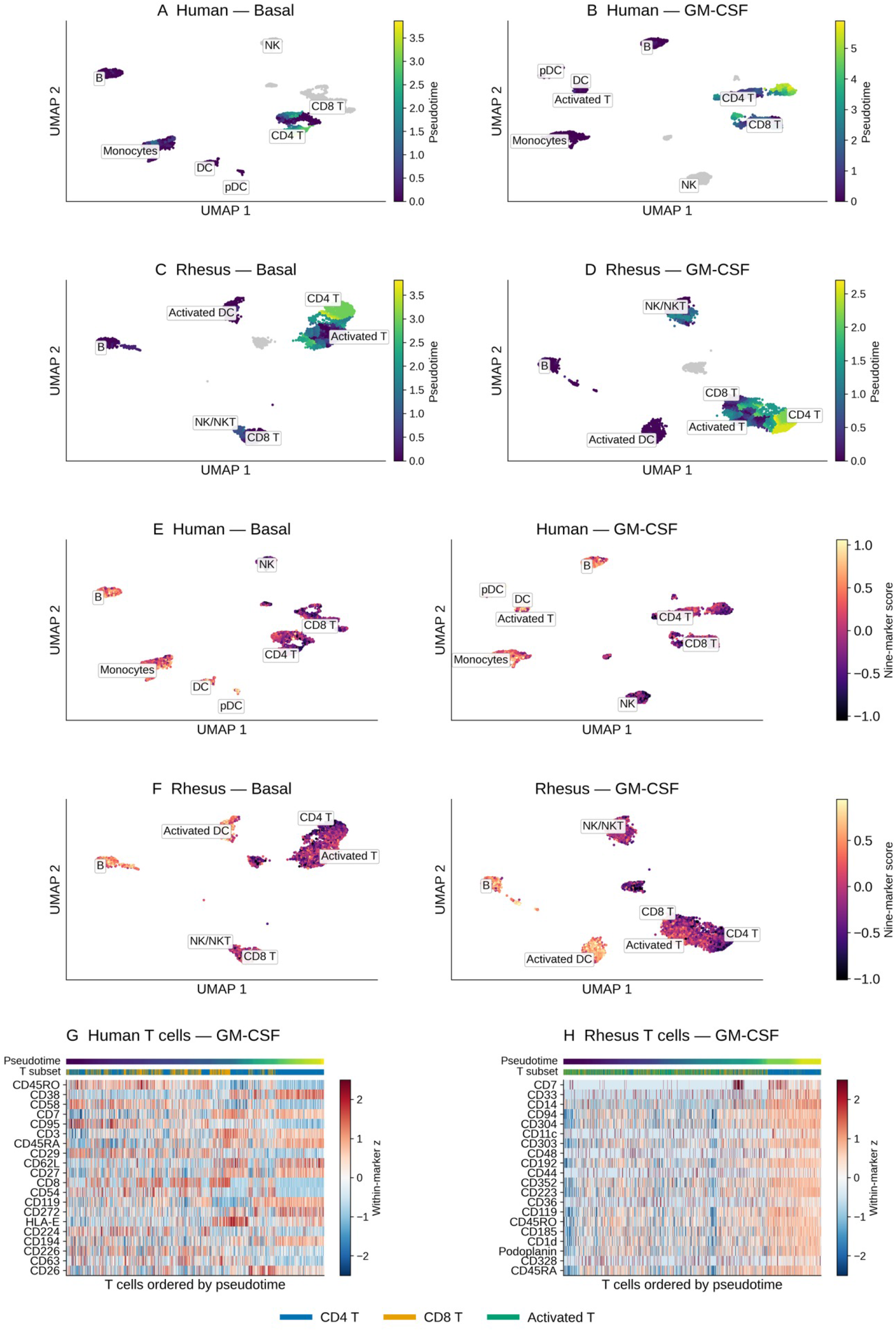
Unstimulated versus GM-CSF population overview and T-cell surface-marker patterns in fused human and rhesus PBMC references. (A–D) Monocle3 pseudotime in human unstimulated (A), human GM-CSF (B), rhesus unstimulated (C) and rhesus GM-CSF (D) fused datasets. CD40L is excluded. QBC specimens are unstimulated; population and condition labels derive from CyTOF partners. Independently fitted graphs use 174 measurements and top-5% nine-marker-score roots; gray denotes nonfinite pseudotime. Pseudotime values are not comparable across graphs. (E, F) Nine-marker proliferation scores for both unstimulated and GM-CSF on the human coordinates in A/B (E) and rhesus coordinates in C/D (F). The score is the equal-weight mean of within-species z-standardized CyTOF Ki67, HLA-DR, pS6, p4EBP1 and pERK1/2, and QBC HLA-DR, CD71, CD25 and CD69, standardized across unstimulated and GM-CSF fused rows only. High scores anchor early pseudotime. Color limits within each species use the combined unstimulated/GM-CSF score 1st/99th percentiles; extreme values are saturated, not excluded. (G, H) Top 20 of 134 QBC markers by absolute Spearman association with finite GM-CSF T-cell pseudotime, in human (G; n = 3,306 fused rows) and rhesus (H; n = 2,987). Existing CD4 T, CD8 T and Activated T labels are retained. Columns follow pseudotime; upper tracks show pseudotime and T subset. Color is within-marker z score across included T-cell-linked rows, saturated at ±2.5. Associations are descriptive, not independent validation.

Among eligible human populations, B cells had the largest absolute mean score change (−0.0363; 23 paired donors). The largest positive mean change was in classical monocytes (+0.0275; 43 donors), narrowly ahead of CD8 T cells (+0.0272; 43 donors). In rhesus macaque, NK/NKT cells had the largest positive and absolute mean change (+0.0635; 12 donors), whereas the activated T population changed by +0.0250 (11 donors). Between-donor SDs for the positive changes in human monocytes, human CD8 T cells and rhesus NK/NKT cells were 0.1272, 0.1190 and 0.0914, respectively. The separate CyTOF-derived pSTAT5 (Y694) feature showed its largest eligible mean increases in human classical monocytes (+0.3411; 43 donors) and rhesus macaque activated dendritic cells (+0.2385; 12 donors), in the existing joint-CLR assay units. These pSTAT5 changes are distinct from the nine-marker score, which does not include pSTAT5.

Given the species-conserved T cell population change with pseudotime (comparing Figure 5A to 5B – human; Comparing Figure 5C to 5D), we were prompted to examined T-cell populations further – utilizing the existing CD4 T, CD8 T and Activated T source annotations/labels. Within each GM-CSF reference graph, QBC markers were ranked by absolute Spearman association with finite pseudotime (Figure 5G, 5H). Human rows (n = 3,306) showed the strongest associations for CD45RO (ρ = −0.491), CD38 (+0.439), CD58 (−0.439) and CD7 (+0.424). In rhesus macaque rows (n = 2,987), CD7 had the strongest association (ρ = +0.518). Positive associations denote higher marker values later in this high-score-rooted ordering; these are descriptive associations among features that also contribute to graph fitting. These patterns describe computationally linked surface phenotypes and reference signaling states along score-rooted, cross-sectional orderings.

### Bayesian network inference reveals convergent pSTAT5 rewiring after GM-CSF stimulation

To ask whether the conserved GM-CSF–pSTAT5 signal sits inside the same network architecture between both species, we inferred condition-specific signaling networks from 15 intracellular measurements^23^, including total IκBα and 14 phospho-readouts, using bnlearn hill-climbing and 500 bootstrap resamples (36 population × condition networks; STAR Methods). Among populations eligible for network analysis, the strongest significant primary GM-CSF pSTAT5 responses occurred in human monocytes and rhesus dendritic cells (Figure 6A–6D). In human monocytes, unstimulated pSTAT5 had majority-oriented inputs from pSTAT1, pS6 and pP38, plus a pMAPKAPK2 connection with unresolved direction. After GM-CSF, the pS6 and pP38 connections were lost; pCREB, pSTAT3, pTBK1/NAK and pERK1/2 connections were gained; the pMAPKAPK2 direction favored pSTAT5→pMAPKAPK2; and the pSTAT1 connection favored pSTAT5→pSTAT1 rather than the unstimulated reverse orientation. In rhesus dendritic cells, the pSTAT1–pSTAT5 connection also reversed its favored direction, pCREB coupling was lost, and pSTAT3 and pERK1/2 coupling was gained. Total retained edge counts changed from 41 to 42 in human monocytes and 25 to 27 in rhesus macaque dendritic cells. These shared local changes support convergent inferred pSTAT5-neighborhood rewiring upon GM-CSF stimulation of human monocytes and rhesus macaque dendritic cells.

**Figure 6.**
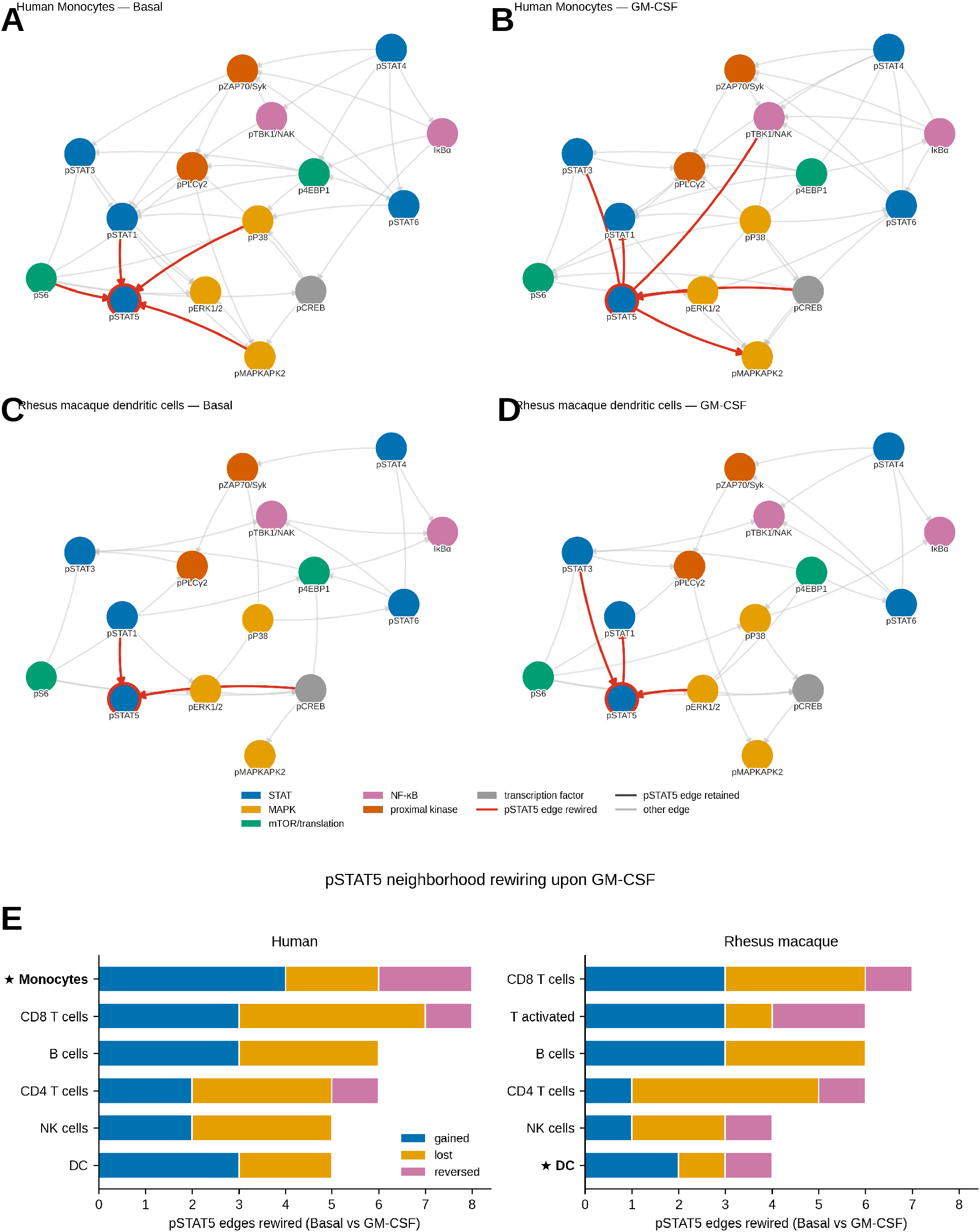
pSTAT5 signaling-neighbourhood rewiring following GM-CSF stimulation in human and rhesus macaque immune cells. (A–D) Majority-oriented bootstrap network summaries of 15 intracellular measurements (14 phospho-readouts and total IκBα) in human monocytes at unstimulated (A) and GM-CSF (B), and rhesus dendritic cells at unstimulated (C) and GM-CSF (D). These were the strongest significant primary GM-CSF pSTAT5 responders among populations eligible for network inference. Networks were fitted separately from primary PBMC-reference fused rows using Gaussian-BIC hill-climbing and 500 row-bootstrap resamples. The archived retained-arc tables implement an 85% support criterion, including boundary values serialized as 0.85. Node positions are fixed across panels. Red edges denote pSTAT5 relationships gained, lost or reversed; dark gray edges denote retained pSTAT5 relationships and light gray other relationships. The pSTAT5 node is outlined in red. Node colors distinguish STAT (blue), MAPK-associated (gold), mTOR/translation-associated (green), NF-κB-associated (pink), proximal signaling (orange) and CREB (gray) measurements. Arrowheads summarize majority bootstrap orientations; approximately equal opposing support denotes unresolved direction. These summaries are not guaranteed to be acyclic and do not establish causal or direct kinase–substrate relationships. Row-bootstrap support is not independent donor-level evidence. (E) Numbers of unique pSTAT5-connected edges gained (blue), lost (gold), or reversed (pink) following GM-CSF stimulation across the indicated human and rhesus macaque populations. Gained and lost edges are present exclusively in the GM-CSF or unstimulated network, respectively; reversed edges are present in both networks but differ in majority bootstrap orientation. Each relationship is counted once in mutually exclusive categories. Stars identify the populations shown in panels A–D.

## Discussion

We combined QBC surface phenotyping with CyTOF intracellular measurements through computational cell matching. The resulting profiles carry 174 measurements from separate cells, providing a framework for describing cross-species immune phenotypes and inferred signaling organization.

First, QBC and CyTOF population frequencies agreed more strongly in human PBMCs (Pearson r = 0.92; n = 9 population–donor pairs) than rhesus macaques (r = 0.55; n = 12 pairs). These are small, correlated collections of population frequencies from three donors per species, not direct measures of single-cell matching accuracy. Under the displayed clustering settings, the modalities resolved complementary populations; the fused resource additionally separated monocyte, dendritic and T-cell subsets and recovered platelet annotations. The 59-marker feature set supports a common fused-clustering space, but antibody specificity, source labels and cross-species resolution remain important constraints.

Second, unstimulated signaling was only partially conserved. B-cell profiles showed the strongest lineage-level conservation (Spearman ρ = 0.70), whereas T-cell, NK and platelet profiles were not significantly correlated across species. Species differences involved total IκBα abundance and selected STAT, TBK1 and CREB phospho-readouts. Total IκBα differences alone cannot determine unstimulated NF-κB activity. The data motivate population-specific comparisons rather than a general rule that innate signaling translates while adaptive signaling does not.

Third, the primary PBMC-reference analysis identified GM-CSF-associated pSTAT5 increases in human monocytes and dendritic populations and rhesus dendritic cells. Cross-species concordance of the complete GM-CSF dendritic-cell response profile was positive but not significant (ρ = 0.511; permutation P = 0.0553), whereas CD40L B-cell and dendritic-cell profiles were significantly concordant. The stronger granulocyte/myeloid GM-CSF correlation belongs to the whole-blood sensitivity analysis and lacks a comparable QBC PBMC counterpart. STAT5 participates in cytokine-dependent immune regulation^24,25,26^, making these observations relevant to future comparative perturbation studies. They do not validate a cross-species pharmacodynamic biomarker or establish therapeutic interchangeability.

Fourth, network summaries showed convergent pSTAT5-centered changes in the featured human monocyte and rhesus dendritic populations: reversal of the favored pSTAT1–pSTAT5 orientation and gained pSTAT3/pERK coupling. These are comparisons of within-condition inferred dependencies, not causal orientations established by interventions. Different featured populations and the targeted panel constrain interpretation. The cell-row bootstrap quantifies stability under that resampling scheme; it does not establish independent donor-level support, correct matching uncertainty or prove that every majority-oriented summary is a directed acyclic graph (DAG). The learned models are also unable to represent cyclic feedback within a single DAG.

The resource can guide more targeted cross-species experiments by identifying populations and signaling nodes for direct testing. Larger donor cohorts, matched native perturbations, dose/time-course measurements and experiments that measure surface and intracellular targets on the same cells would be needed to establish the biological and translational scope of the inferred patterns. Published variation in human cytokine responses further motivates donor-aware validation^27^.

Several limitations qualify these conclusions. The QBC measurements derive from PBMCs, whereas the published stimulation reference derives from whole blood. Restricting the primary matching reference improves compositional compatibility but cannot ensure that every computational pair represents an equivalent cell state. Granulocyte/myeloid results are therefore kept in the separate whole-blood sensitivity analysis. Rhesus T-cell subset resolution is also limited by the shared marker panel and reagent cross-reactivity; fine T-subset labels should be interpreted cautiously.

MARIO links separately measured cells, potentially across donors; the fused features are not physical same-cell measurements. QBC provides an unstimulated surface reference, while stimulation and analysis population labels come from CyTOF partners. Reused reference cells and shared QBC inputs mean that fused rows are not independent biological replicates. Reference-cell-first donor summaries mitigate reuse in the Figure 5 comparisons but do not capture all matching or QBC-donor uncertainty; network bootstraps resample fused rows rather than independent donors. Pseudotime is cross-sectional and deliberately score-rooted, so score–pseudotime association is not independent temporal validation. QBC marker associations are descriptive because those features also enter graph fitting. T-subset composition, source annotation and antibody-specificity can influence these trends.

## STAR Methods

### Resource Availability

Lead contact: Further information and requests for resources and reagents should be directed to and will be fulfilled by the lead contact, David R. McIlwain.

Materials availability: Antibody panels and the shared-marker definitions are supplied in Tables S1–S4 and the Key Resources Table. The QBC method is summarized in STAR Methods and Figure S1. Requests for custom-conjugated reagents should be directed to the lead contact and may be subject to a materials transfer agreement.

Data and code availability: Raw data deposited for this study are available at Zenodo (https://doi.org/10.5281/zenodo.23676187). Analysis code and processed objects are available in the public GitHub repository (https://github.com/aaron-tantk/QBC-CyTOF_cross-species_PBMCs); the manuscript-aligned release is v1.1.0 (https://github.com/aaron-tantk/QBC-CyTOF_cross-species_PBMCs/releases/tag/v1.1.0). Tables S1–S6 accompany this submission. Additional information required to reanalyze the data is available from the lead contact upon request.

### Experimental Model and Subject Details

Human PBMCs: Human peripheral blood mononuclear cells were isolated by density-gradient centrifugation on Ficoll-Paque. Human samples were collected with written informed consent under Stanford University IRB protocol #28289. Demographics for the three experimental donors (H1–H3) are summarized in Table S6. PBMCs were cryopreserved in 90% FBS/10% DMSO and stored at −80 °C until use.

Rhesus macaque PBMCs: Three rhesus macaque (Macaca mulatta) cryopreserved PBMC samples (M1– M3) were obtained from CellCarta. Animal sample collection was conducted under Stanford APLAC #26675 and followed the NIH Guide for the Care and Use of Laboratory Animals. Sample-source information, including source-confirmed sex and dates of birth, is listed in Table S6; age at collection is not reported.

Published cross-species CyTOF stimulation dataset: The whole-blood stimulation reference CyTOF dataset of human and rhesus macaque samples was previously generated and described^18^. For this study, the unstimulated, CD40L-stimulated, and GM-CSF-stimulated conditions were selected from that dataset’s 15-stimulus panel (46 human and 15 rhesus donors; raw export capped at 4,000 cells per donor per condition), balanced to 260 (human) or 800 (rhesus macaque) cells per donor per condition and arcsinh-transformed (cofactor 5), for cross-modal MARIO fusion with our cross-species QBC reference. After PBMC-lineage restriction of the reference, matching, and fusion quality control, the primary fused stimulation objects contained 20,862 human and 14,414 rhesus cells (human: 7,740 unstimulated / 6,164 CD40L / 6,958 GM-CSF; rhesus: 4,807 / 4,618 / 4,989); fusion against the full whole-blood reference (20,550 human and 13,501 rhesus cells) was retained as a sensitivity analysis.

### Method Details

Quantum Barcoding (QBC) workflow: Single-cell protein expression was profiled using a workflow adapted from the published QBC method^14^ with the study-specific conditions below. Cells were fixed with 1.6% paraformaldehyde for 10 min at room temperature, pelleted (600 × g, 3 min, 4 °C), and permeabilized on ice for 10 min using ice-cold methanol. Fixed/permeabilized cells were stored in cell-staining medium (0.5% BSA in PBS) at –80 °C until use. Cryopreserved fixed cells were rehydrated through sequential washes in SME buffer (1× PBS, 0.5% BSA, 0.02% sodium azide, 5 mM EDTA) and HSME buffer (SME containing 0.5 M NaCl). Non-specific binding was blocked by incubation in 0.5 M NaCl, TruStain FcX, and 300 ng/µL salmon sperm DNA in SME buffer for 10 min on ice. For staining, 2 × 10⁶ cells per sample were incubated with pre-titrated pools of oligo-conjugated antibodies (TotalSeq-B Universal Panel v1.0; BioLegend) for 30 min at room temperature. Stained cells were re-fixed with 1.6% paraformaldehyde for 10 min at room temperature to crosslink antibody–oligo complexes, washed twice with HSME buffer, counted, and proceeded to combinatorial barcoding. Four rounds of 60-way split-and-pool barcoding were performed; cells were distributed among 60 aliquots per round, ligated to an index oligonucleotide with the corresponding block solution, and pooled between rounds. After the fourth round, library preparation used 50,000 cells per PCR. Cells were lysed and RNA degraded as follows: 50,000 cells were combined with 5 U RNase H (NEB M0297), 1× RNase H reaction buffer, and 10 mM DTT in 20 µL, incubated at 37 °C for 30 min, followed by 1.4 mg/mL Proteinase K (NEB P8107S) at 55°C for 1 h, then 95 °C for 10 min to inactivate. Second-strand cDNA synthesis and PCR were performed in 50 µL with 0.4 mM dNTPs, 0.025 µM RNA-specific PCR primers, 0.5 µL Herculase II Fusion DNA Polymerase (Agilent 600675), and 1× Herculase II reaction buffer; cycling: 95 °C / 2 min initial denaturation followed by 1–20 cycles of 95 °C / 20 s, 62 °C / 20 s, 68 °C / 1 min, and 68 °C / 4 min final extension. PCR products were purified on SPRI beads (1:1), quantified by Qubit hsDNA assay, and sized on Agilent Bioanalyzer or TapeStation.

CyTOF data acquisition: CyTOF analysis of cryopreserved human and rhesus macaque PBMCs followed our previously described protocol^18^. Briefly, PBMCs were thawed in Cell Staining Medium (PBS + 0.5% BSA + 0.02% sodium azide + 25 U/mL benzonase) and fixed in 1.6% paraformaldehyde for 10 min. Fixed cells were incubated 45 min at room temperature with a cocktail of metal-tagged surface antibodies for cross-species immunophenotyping, then permeabilized by dropwise addition of ice-cold 100% methanol with gentle vortexing and incubated on ice for 10 min. Cells were washed and stained for 60 min with a panel of metal-tagged intracellular antibodies, resuspended in iridium intercalator solution (Standard Biotools) containing 1.6% paraformaldehyde, and acquired on a freshly tuned CyTOF2 mass cytometer (Standard Biotools). Data were normalized using bead standards and analyzed in CellEngine (CellCarta) and custom R scripts.

Single-modality and unstimulated fused analyses: CyTOF FCS intensities were arcsinh-transformed with cofactor 5, imported into Seurat and CLR-normalized (margin = 2). Figure 3A/3B and Figure S2A/S2B contain 36,000 CyTOF cells per species: 12,000 from each donor. QBC count tables were CLR-normalized (margin = 2) and donor/site datasets integrated using FindIntegrationAnchors and IntegrateData^29^. Figure 3C/3D and Figure S2C/S2D contain 40,000 QBC cells per species: 10,000 from each donor at Stanford, plus an additional 10,000-cell Davis dataset from H1 in human and M3 in rhesus. These are additional site datasets, not additional donors. The Stanford-only QBC matching export contained 30,000 cells per species. The saved standalone CyTOF PCA used 25 human or 24 rhesus non-phospho/context features, with neighbors and UMAP on PCs 1–16; QBC PCA used 134 human or 118 rhesus features, with neighbors and UMAP on PCs 1–10. Selected standalone cluster assignments correspond to resolution 0.3, consolidated into 10/9 CyTOF and 7/6 QBC annotated populations in human/rhesus. The unstimulated fused resource retained 29,412 human and 28,578 rhesus pairs. Its separate clustering used 59 QBC markers, PCs 1–20, cosine UMAP and recorded Louvain resolution 0.7. The final saved annotation map consolidates 18 human and 13 rhesus graph clusters into 14 human and 11 rhesus annotated populations.

MARIO cross-modal integration: QBC and CyTOF datasets were integrated using MARIO^20^. The matching was anchored on 18 surface markers shared between the QBC and CyTOF panels (Table S3), including the lineage-defining markers CD3, CD11b, CD11c, CD14, CD19 (CD20 in rhesus macaque), CD56, and HLA-DR. For the unstimulated matching resource, per-species inputs were balanced to 30,000 cells per modality by random subsampling (random_seed = 42) prior to matching. The MARIO pipeline (pipelined_mario) was run with normalization enabled on 5 batches per modality: an initial distance matrix on the 18 shared features (10 canonical components), one-to-one matching (n_matched_per_cell = 1, sparsity = 400), a refined all-feature distance matrix (12 canonical components), interpolation of matching weights (4 weights, 5 canonical-correlation components), and statistical filtering of low-confidence matches (10 clusters, 10 components, bad_prop = 0.2, 20 iterations), followed by joint embedding by canonical correlation analysis (10 dimensions, 500 iterations). CyTOF-derived intracellular signaling features were then concatenated onto the matched QBC cells to produce the 174-feature integrated dataset per species. Integration was performed independently for the unstimulated cross-species resource and for the stimulated cross-modal fusion described below. For the stimulation analysis, MARIO was applied to fuse the published CyTOF whole-blood stimulation dataset^18^ with this study’s QBC cross-species surface-ome (unstimulated, 3 donors per species, per-donor CLR-normalized). For the primary stimulation analyses, CyTOF reference cells annotated as granulocyte/myeloid, platelet, erythrocyte, or unresolved by de novo clustering of the balanced reference were excluded before matching (PBMC-only design; 13,376 human and 24,684 rhesus macaque reference cells), yielding the 20,862-cell human and 14,414-cell rhesus fused stimulation objects described above; matching against the full whole-blood reference (20,550 human and 13,501 rhesus macaque pairs) was retained as a sensitivity analysis. The underlying biological assumption (that short-term ligand stimulation primarily modulates intracellular signaling without changing surface phenotype) is consistent with prior time-course profiling of these stimuli^18^.

Matching validation: Match quality was assessed by (i) within-pair Pearson correlation across the 18 shared anchor markers, compared against a shuffled-partner null distribution; (ii) Harmony integration^21^ of the two modalities on the 18 anchors (per-modality z-scoring, PCA, RunHarmony with modality as the grouping variable) with 1-nearest-neighbor label transfer from CyTOF to QBC cells; and (iii) CellFuse (preprint) supervised contrastive learning^22^ trained on the annotated CyTOF reference (70/30 stratified train/validation split; learning rate 9×10^−4^;, margin 0.8, batch size 256, up to 50 epochs with early stopping, 8-dimensional output embedding, leaky-ReLU, dropout 0.7) with 5-nearest-neighbor label transfer to QBC cells. Cross-method agreement was quantified on the matched pairs by percent concordance, adjusted Rand index, and normalized mutual information. Stimulation-condition mixing of the fused embedding was quantified by the condition local inverse Simpson index (cLISI; 30 nearest neighbors) on the joint MARIO embedding.

Stimulation fused clustering and analysis labels: The matched profiles carried 174 features (134 QBC and 40 CyTOF), with modality suffixes. The joint assay was CLR-normalized within each fused row (margin = 2). Clustering used the 59-marker QBC subset, 20 PCs, FindNeighbors on PCs 1–20 and FindClusters at resolution 0.5, with resolution 1.2 as a finer sensitivity analysis. The primary resolution-0.5 solution contained 16 human and 11 rhesus graph clusters. Figures 4 and 6 retained broad CyTOF source-cell population labels, while Figure 5 retained fine CyTOF reference-cell labels; these results were not generated by collapsing cluster-plurality labels. Cluster-plurality assignments, compositions, purity and within-pair anchor correlations are supplied only as an annotation/match-quality audit in Table S5, with primary and sensitivity designs explicitly separated.

Nine-marker signature and population summaries: The nine-marker proliferation signature comprised CyTOF Ki67, HLA-DR, pS6 (S235/S236), p4EBP1 (T37/T46) and pERK1/2 (T202/Y204), together with QBC HLA-DR, CD71, CD25 and CD69. Each measurement in the existing joint-CLR assay was standardized across all included fused rows within species, including only unstimulated and GM-CSF reference conditions; the nine z scores were then averaged with equal weights. The QBC HLA-DR assay feature was HLA-DR-QBC, resolving the HLA.DR-QBC naming alias, and all nine nonconstant features were required. For population comparisons, scores were first averaged across repeated links to each unique CyTOF reference cell and then across unique reference cells within reference donor, population and condition group. CD40L was excluded from this comparison. GM-CSF-minus-unstimulated differences were paired within reference donor; a donor required at least three unique reference cells in each group, and a population required at least five eligible paired donors for ranking by the absolute mean paired difference; the largest positive mean difference was also identified. Direction, donor counts and between-donor SDs were reported descriptively, without new population-level significance tests. The targeted CyTOF-derived pSTAT5 (Y694) comparison used the same reference-cell averaging and paired-donor criteria on the existing joint-CLR assay values, without feature z-standardization; pSTAT5 was not included in the nine-marker signature.

Bayesian network inference: We applied structure learning to primary PBMC-reference fused profiles^23^. Groups were defined by the CyTOF partner’s broad source label, not by plurality transfer from fused clusters. Eligible human groups were B cells, CD4 T cells, CD8 T cells, DCs, monocytes and NK cells; eligible rhesus groups were B cells, CD4 T cells, CD8 T cells, DCs, NK cells and activated T cells. Each was split into Unstimulated, CD40L and GM-CSF groups, requiring at least 100 fused rows per condition. Input variables were joint-CLR assay values for pSTAT1 (Y701), pSTAT3 (Y705), total IκBα (amino-terminal), p4EBP1 (T37/T46), pZAP70/Syk (Y319/Y352), pS6 (S235/S236), pCREB (S133), pSTAT4 (Y693), pSTAT5 (Y694), pTBK1/NAK (S172), pSTAT6 (Y691), pPLCγ2 (Y579), pERK1/2 (T202/Y204), pP38-MAPK (T180/Y182) and pMAPKAPK2 (T334). Zero-variance features were excluded. The bnlearn package^30^ performed hill-climbing with Gaussian BIC (hc, score = “bic-g”) and 500 bootstrap resamples of fused rows. An 85% bootstrap-support criterion was used; displayed results reuse archived retained-arc tables, including boundary values serialized as 0.85. For summary/plotting, the two direction rows for each unordered variable pair were collapsed to one edge using the majority bootstrap direction; unresolved directions were described as such. The summaries contain 36 population × condition networks. pSTAT5 rewiring counts unique connected edges gained, lost or reversed between unstimulated and GM-CSF, counting a reversal once. Featured human monocyte and rhesus DC groups were selected from eligible strong GM-CSF pSTAT5 responders; this is a descriptive selection, not a prospectively validated criterion. The displayed majority-oriented summaries are not automatically equivalent to bnlearn’s averaged acyclic networks. Directional support does not resolve causal identifiability or establish direct biochemical regulation.

### Quantification and Statistical Analysis

Cell population quantification: QBC reads were processed into cell-barcode, UMI and antibody-tag counts. The Figure 2A/2B proof-of-concept run used the UMI-rank knee to select a minimum of 1,000 UMIs. Figure 2 population frequencies were calculated separately within donor using the manual-gating denominators specified in its legend; these frequency estimates are distinct from Figure 3 clustering displays. No uniform 10,000-cell sampling assumption is imposed across Figures 2 and 3. Sample origin in the cross-species workflow was assigned through SC-A sample-index ligation, not post-hoc hashtag deconvolution. Standalone and fused feature spaces, clustering resolutions and display counts are described above; UMAP was used for visualization.

Cross-platform concordance: QBC vs. CyTOF concordance for population frequencies was assessed on canonical lineage labels harmonized across platforms (human: B cells, T cells, classical monocytes; rhesus macaque: B cells, dendritic cells, myeloid cells, T cells), each measured in three donors per species (n = 9 and 12 population–donor pairs, respectively), using two-tailed Pearson correlation and simple linear regression. Population–donor points share donors and are constrained by composition. Reported P values and regression confidence bands are nominal point-level summaries, not donor-aware inference; there are only three biological donors per species.

Stimulation-response statistical analysis: Stimulation responses were quantified as donor-mean differences (stimulated − unstimulated) per population and intracellular signaling measurement, using the CyTOF-side values of fused pairs. Within-species significance was assessed by paired, two-tailed t-tests across donors (combinations with fewer than 5 donors, or fewer than 3 cells per donor per condition, were not tested), with Benjamini–Hochberg FDR across all population × node × condition tests (375 tests in the primary PBMC-only analysis; 480 in the whole-blood sensitivity analysis). Cross-species conservation of the 15-node response profiles was assessed by Spearman correlation of donor-mean response vectors within harmonized lineages and globally, with two-sided stratified permutation tests (10,000 permutations, permuting node correspondence within lineage strata).

Bayesian network quantification: At least 100 fused rows per condition were required for each included population. There were 500 row-bootstrap iterations and an 85% support criterion, with 36 archived population × condition outputs. Edge and rewiring summaries collapse duplicate direction rows into unique pairs as described above; reversals count once. Bootstrap support is not a donor-level P value.

Pseudotime analysis: Four trajectories were fitted in Monocle3^31^ (version 1.3.1; igraph 1.6.0), separately for human and rhesus macaque unstimulated and GM-CSF reference groups, with CD40L excluded. The existing CLR-normalized Protein assay included all 174 fused measurements; metadata were excluded. We used preprocess_cds(num_dim = 50, norm_method = “none”), cosine UMAP with 15 neighbors and min_dist = 0.1, cluster_cells and learn_graph(use_partition = TRUE, close_loop = TRUE), with random seed 42, including the PCA and UMAP stages. Root cells were those at or above the within-graph 95th percentile of the nine-marker score; order_cells assigned pseudotime, with disconnected/unrooted components left nonfinite. Finite pseudotime covered 5,250/7,740 human unstimulated, 5,920/6,958 human GM-CSF, 4,507/4,807 rhesus unstimulated and 4,646/4,989 rhesus GM-CSF fused rows. For Figure 5G and 5H, T-cell-linked rows retained the existing CD4_T_cells, CD8_T_cells and Activated_T_cells labels. All 134 QBC markers were ranked by absolute Spearman correlation with finite GM-CSF T-cell pseudotime. The top 20 per species were plotted in pseudotime order using within-marker z scores across the displayed T-cell-linked rows, with display colors saturated at ±2.5. These rankings were descriptive; no Moran’s I significance filter was applied.

Software: FCS input used flowCore^32^. Normalization, integration, clustering and marker analysis used Seurat^29,33^. Trajectories used Monocle3^31^ version 1.3.1 with igraph 1.6.0; Bayesian network learning used bnlearn^30^. Network layouts used igraph, tidygraph and ggraph. R visualization used ComplexHeatmap^34^, pheatmap, ggplot2^35^ and RColorBrewer. Python processing used MARIO (mario-py), NumPy, pandas, scikit-learn, SciPy, matplotlib and adjustText; gating used CellEngine (CellCarta). Stimulation-response and marker-association plots were generated programmatically from the saved data. Generative image tools were used only for the explanatory workflow schematics identified in their legends, not to synthesize or alter data panels.

### Key Resources Table

| REAGENT or RESOURCE | SOURCE | IDENTIFIER |
|---|---|---|
| **Antibodies** |  |  |
| TotalSeq-B Universal Panel v1.0 (134 antigen-targeting reagents plus six isotype controls) | BioLegend | Cat. #199902; Table S2 |
| Human CyTOF surface panel (within the 42-marker study panel; Table S1) | This study; workflow adapted from published profiling<sup>18,19</sup> |  |
| Rhesus CyTOF surface panel (within the 43-marker study panel; Table S1) | This study; workflow adapted from published profiling<sup>18,19</sup> |  |
| CyTOF intracellular signaling/proliferation panel (includes total IκBα and Ki67; not all phospho-antibodies; Table S1) | This study; workflow adapted from published profiling<sup>18,19</sup> |  |
| TruStain FcX Fc receptor blocker | BioLegend | Cat. # 422301 |
| **Biological Samples** |  |  |
| Human PBMCs (n = 3 donors: H1, H2, H3) | Consented Donors of Nolan Laboratory, Stanford University IRB protocol #28289 |  |
| *rhesus macaque* PBMCs (n = 3 animals: M1, M2, M3) | CellCarta Fremont LLC (Fremont, CA) and Stanford University APLAC #26675 |  |
| **Chemicals, peptides, and recombinant proteins** |  |  |
| Recombinant human GM-CSF | Cell Sciences | Cat. # CRG100B |
| Recombinant human/ *rhesus macaque* CD40L | Enzo Life Sciences | Cat. # ALX-522-110-C010 |
| Paraformaldehyde 16% | Thermo Fisher Scientific | Cat. # 28906 |
| Methanol (HPLC grade) | Sigma-Aldrich | Cat. # 34860 |
| Bovine Serum Albumin (BSA) | Sigma-Aldrich | Cat. # A7906 |
| RNase H | New England Biolabs | M0297 |
| Proteinase K | New England Biolabs | P8107S |
| Herculase II Fusion DNA Polymerase | Agilent | 600675 |
| Salmon Sperm DNA | Thermo Fisher Scientific | Cat. # AM9680 |
| **Commercial Assays** |  |  |
| Legacy Scale QBC Protein workflow (TotalSeq-B antibody chemistry) | Scale Biosciences; supplied undated protocol<sup>28</sup> | Study-specific conditions in STAR Methods and Figure S1; not the newer TotalSeq-A PhenoCyte workflow |
| Maxpar X8 Antibody Labeling Kit | Standard Biotools | Cat #: 201300 |
| Qubit dsDNA HS Assay Kit | Thermo Fisher Scientific | Q32854 |
| Agilent 2100 Bioanalyzer / TapeStation High Sensitivity DNA Kit | Agilent Technologies |  |
| **Deposited Data** |  |  |
| Public raw QBC and CyTOF data | This study | Zenodo: 10.5281/zenodo.23676187 |
| Revised MARIO-integrated 174-feature objects and derived analysis packages | This study | GitHub release v1.1.0: https://github.com/aaron-tantk/QBC-CyTOF_cross-species_PBMCs/releases/tag/v1.1.0; raw data at Zenodo: 10.5281/zenodo.23676187 |
| Published cross-species CyTOF stimulation reference | Bjornson-Hooper et al.<sup>18</sup> | FlowRepository: FR-FCM-Z5PC |
| **Software and Algorithms** |  |  |
| R | R Foundation | 4.4.2 |
| bnlearn (network structure learning) | Scutari, 2010<sup>30</sup> | https://www.bnlearn.com |
| Monocle3 1.3.1 (pseudotime) | Cao et al., 2019<sup>31</sup> | https://cole-trapnell-lab.github.io/monocle3/ |
| igraph, tidygraph and ggraph (network layouts) | Open-source R packages | https://r.igraph.org; https://tidygraph.data-imaginist.com; https://ggraph.data-imaginist.com |
| Seurat (normalization, integration and clustering) | Hao et al., 2024; Stuart et al., 2019<sup>29,33</sup> | https://satijalab.org/seurat/ |
| flowCore (FCS handling) | Hahne et al., 2009<sup>32</sup> | https://bioconductor.org/packages/flowCore/ |
| MARIO (cross-modal cell matching) | Zhu et al., 2023<sup>20</sup> | https://github.com/shuxiaoc/mario-py |
| CellEngine (CyTOF gating) | CellCarta | https://cellengine.com |
| Custom QBC processing and analysis scripts | This study | https://github.com/aaron-tantk/QBC-CyTOF_cross-species_PBMCs |

## Acknowledgments

The authors thank previous members of Scale Biosciences Inc. for their unwavering technical support. The authors thank members of the Nolan and McIlwain laboratories for helpful discussions and technical assistance. The authors thank Dr. Shigemi Kinoshita for technical assistance.

## Author Contributions

T.K.T. and D.R.M. conceived of the idea, designed and performed experiments, analyzed data, and wrote the manuscript. K.T., F.S., F.Z., B.Z., Y.B., S.A., D.L.L.H., Z.B.-H., M.A.B., and A.T. contributed tools, reagents, and experimental materials. B.Z. developed the software used in data analysis. K.T. provided experimental materials, reagents, and software in data analysis. H.N., A.C., G.P.N., and D.R.M. supervised the project, reviewed, and edited the manuscript.

## Declaration of generative AI and AI-assisted technologies in the manuscript preparation process

During the preparation of this work, the authors used Claude (Anthropic) for language editing and to check the manuscript for clarity, consistency, and formatting. The authors reviewed and edited the output as needed and take full responsibility for the content of the published article.

## Figure and Figure Legends

**Supplementary Figure 1:**
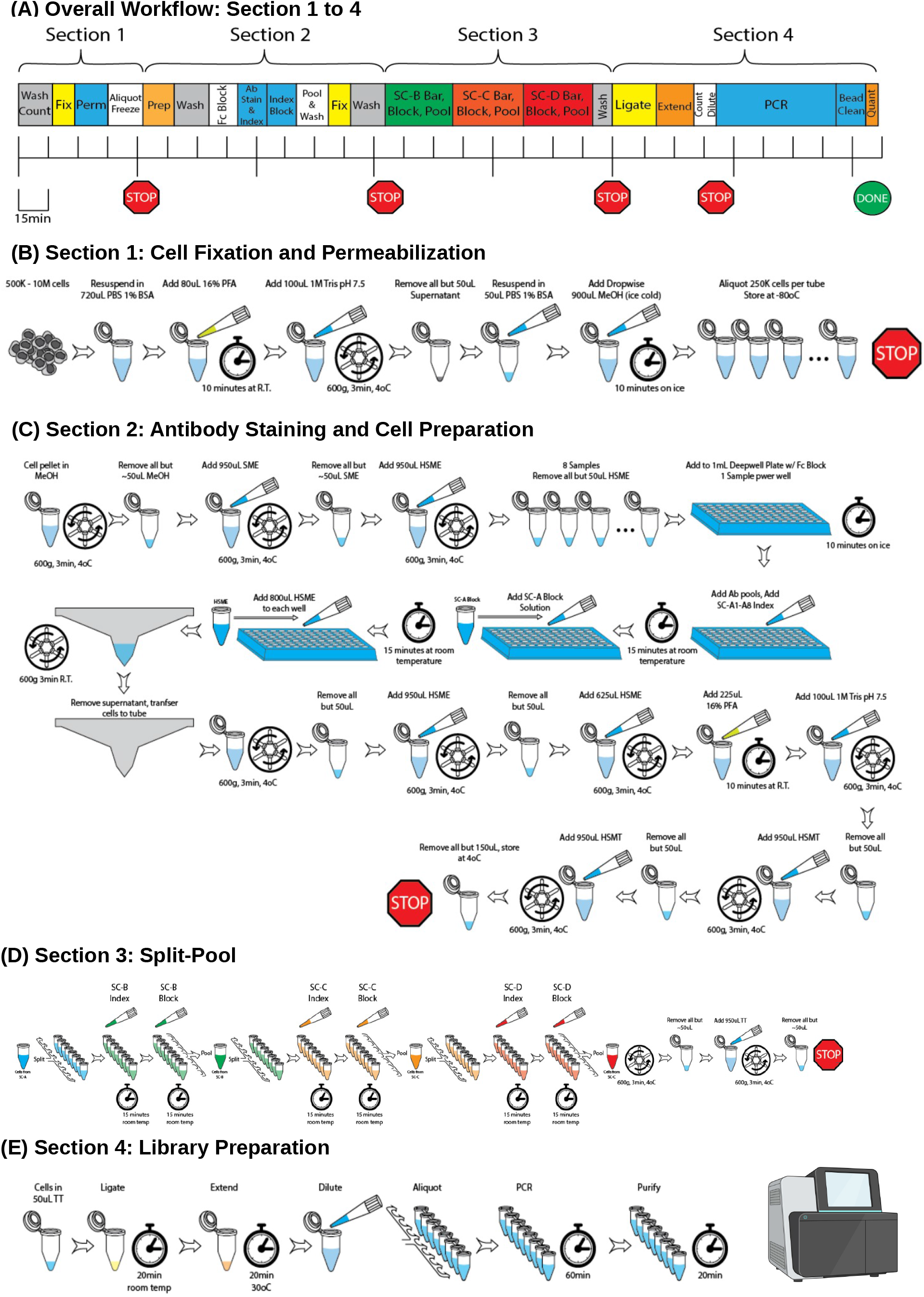
Detailed workflow of the method for single-cell profiling of protein expression through next generation sequencing. Antibody–oligonucleotide conjugates encode protein targets, and combinatorial DNA indices support cell-level assignment of sequencing counts. The legacy illustration contains generic quantities and an eight-way indexing scheme that do not represent the study-specific conditions below; see STAR Methods for the adapted QBC workflow. (A) Workflow overview: Cell fixation and permeabilization, antibody staining, four 60-way split-pool barcode rounds, and sequencing-library preparation. (B) Cell fixation and permeabilization: Cells were initially fixed with 1.6% paraformaldehyde for 10 min at room temperature and permeabilized with ice-cold methanol on ice for 10 min. (C) Antibody staining and cell preparation: 2 × 10⁶ cells per sample were stained with oligo-conjugated antibodies for 30 min at room temperature. After staining, cells were re-fixed with 1.6% paraformaldehyde for 10 min at room temperature and washed before combinatorial barcoding. (D) Split-pool barcoding: Four 60-way rounds were performed, with index ligation in each round and pooling between rounds. The nominal combinatorial capacity is 60^4^ = 12,960,000; this is not a measured singlet yield or collision rate. (E) Library preparation: Barcoded cells were processed using 50,000 cells per PCR. Amplified products were purified with SPRI beads, quantified using the Qubit hsDNA assay, and sized using an Agilent Bioanalyzer or TapeStation before sequencing.

**Figure S2.**
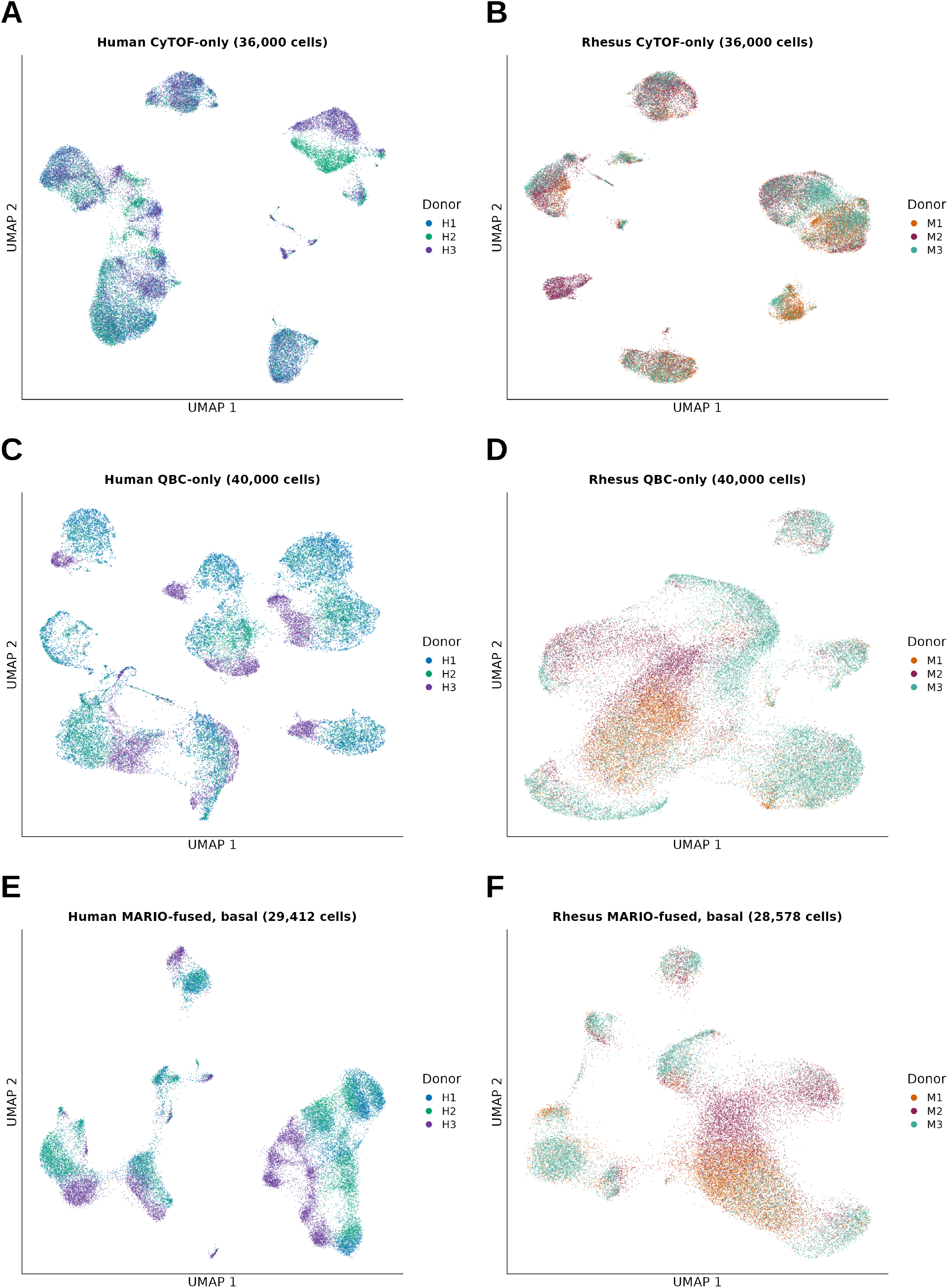
Donor-colored embeddings of standalone and unstimulated fused datasets. (A–F) UMAP embeddings corresponding to Figure 3A–3F, colored by donor: human H1–H3 and rhesus M1–M3. (A, B) CyTOF-only; (C, D) QBC-only; (E, F) unstimulated MARIO-fused profiles colored by QBC-side donor. Standalone QBC includes unequal donor/site sampling as described in Figure 3. MARIO matching was donor-agnostic: QBC/CyTOF donor concordance was 49% in human and 35% in rhesus pairs. Donor-colored embeddings are descriptive and do not establish absence of donor or batch effects.

